# Adipose-derived Mesenchymal Stem Cells and Retinal Pigment Epithelial Cells Interactions in Stress Environment via Tunneling Nanotubes

**DOI:** 10.1101/2024.11.24.624852

**Authors:** Merve Gozel, Karya Senkoylu, Cem Kesim, Murat Hasanreisoglu

## Abstract

This study aims to demonstrate the formation of TNTs between AdMSCs and RPE-1 and their alterations in response to experimental stress conditions. Serum starvation was employed as a stress condition to induce TNTs between the AdMSC and RPE-1. The presence of TNTs was demonstrated through immunofluorescence microscopy while scanning electron microscopy was utilized to determine the average thickness. Cell viabilities were assessed after stress by CTG, and H2DCFH-DA probes evaluated the cells’ reactive oxygen species (ROS) levels. Further, JC-1 labeled mitochondrial exchange between cells via TNTs was supported by videos. A transmembrane culture system was employed to inhibit TNT formation. In this study, we investigated the role of TNTs in facilitating intercellular communication and mitochondrial transfer between AdMSCs and RPE-1 under stress. We found that TNT-mediated mitochondrial transfer from AdMSCs to RPE-1 helps to reduce ROS levels and improve cell viability. We demonstrated that direct interaction between AdMSCs and RPE-1 was crucial for stress recovery. Co-culture enhanced viability and sustained retinal epithelial cell function after stress-induced damage. Mechanical inhibition of TNT formation decreased cell viability and increased ROS levels, indicating the importance of TNTs in cellular protection. The findings can provide a new perspective on the therapeutic potential of stem cell-based therapy in protecting RPE against stress-induced damage and promoting tissue regeneration.

## 1. Introduction

TNTs are long cytoplasmic bridges that have a remarkable ability to conduct a wide range of functions that mediate cellular physiology and cell survival (Tiwari, Koganti et al. 2021). TNT appears to play a role in intercellular exchanges of signals, chemicals, organelles, and pathogens, involving them in a wide range of functions, according to a growing body of research (Dupont, Souriant et al. 2018). They are crucial in embryonic development, cell migration, cell healing, cancer therapy resistance, and pathogen dissemination as intercellular bridges (Dupont, Souriant et al. 2018). The presence of TNTs are confirmed in various ocular cell groups including corneal (Chinnery and Keller 2020), trabecular (Quigley 2011), and retinal (Wittig, Wang et al. 2012) cells. The retinal pigment epithelium (RPE) is a monolayer of cells located at the posterior of the eye, between the retina and choroid, which has critical roles in vision (Yang, Zhou et al. 2021). Damage to the structure and function of the retinal pigment epithelium leads to a variety of retinal diseases such as diabetic retinopathy, age macular degeneration, or retinitis pigmentosa. Novel treatment strategies for restoring the structural and functional integrity of RPE cells lead to the ocular application of mesenchymal stem cells (MSCs). MSCs have been shown to have beneficial effects on damaged RPE cells. These effects include the differentiation of MSCs into RPE cells (Zhu, Chen et al. 2022) and the regulation of the retinal medium through MSC-derived conditioned media (Eiro, Sendon-Lago et al. 2022). Adipose-derived MSCs (AdMSCs) have been previously shown to establish TNTs directed to corneal epithelial cells to perform mitochondrial transfer aiming for protection against mitochondrial damage (Jiang, Gao et al. 2016). AdMSCs are particularly advantageous due to their easy accessibility, high yield, and fewer ethical concerns compared to other stem cell sources (Shi et al. 2024). Given that various cell types of the ocular tissue including RPE cells show the presence of TNT structures, and there is increasing evidence for MSCs to have TNTs as another mechanism of action aimed for cell and tissue regeneration, the question arises whether a similar mechanism will also be employed in retina pigment epithelium repair following retinal injury.

In this study, we aimed to determine whether AdMSC-mediated mitochondrial transfer could rescue the retinal pigment epithelium from several stress-induced mitochondrial damages. We established that AdMSCs could efficiently donate functional mitochondria and protect RPE-1 from serum starvation stress-induced damage through TNT formation.

## 2. Materials and Methods

### 2.1 Ethics Statements

This study adhered to national regulations and was approved by the Ethics Committee of the Koc University School of Medicine, Istanbul, Turkey (protocol no: 2020.447.IRB2.121, date of approval : 03.12.2020). Study subjects (8 adipose tissues from liposuction surgery) were recruited from Koç University Hospital, Plastic, Reconstructive and Aesthetic Surgery Department. Written informed consent was obtained from all patients after an explanation of the nature and possible consequences of the study.

### 2.2 Isolation and culture of adipose-derived mesenchymal stem cells (AdMSCs)

Adipose-derived mesenchymal stem cells (AdMSCs) were isolated, cultured, and characterized as previously described (Gozel et al. 2024). Briefly, AdMSCs were isolated from liposuction samples obtained from healthy male and female donors (aged 20–50) and processed using collagenase digestion. Cells were cultured in Dulbecco’s Modified Eagle Medium - low glucose (DMEM-LG; Biowest, Nuaille, France) supplemented with 10% (v/v) Fetal Bovine Serum (FBS; Biowest, Nuaille, France) and passaged until P4 for co-culture experiments. Characterization of AdMSCs was performed via flow cytometry using positive markers (CD73, CD90, CD146) and negative markers for hematopoietic cells (CD3/4/8, CD56, CD20). Differentiation potential into osteogenic, chondrogenic, and adipogenic lineages was assessed using commercial differentiation kits, with lineage-specific markers (Alizarin Red S, Alcian Blue, and Oil Red O) confirming differentiation through microscopy imaging.

### 2.3 Culturing of Retinal Pigment Epithelial Cell Line (RPE-1)

RPE-1 cells (ATCC®□ CRL□4000) were grown in DMEM/F12 (Dulbecco’s MEM; PAN Biotech, Aidenbach, Germany) supplemented with 10% fetal bovine serum (Biochrom, Berlin, Germany), 100 IU/ ml penicillin, 100 μg/ml streptomycin and were used at passages 10 to 15. The viability of cultured cells was assessed microscopically by the trypan blue exclusion test.

### 2.4 In vitro co-culture systems

#### 2.4.1 Oxidative stress model and in vitro co-culture system

RPE-1 with (oxidative stress) or without (normal condition) hypoxia treatment were co-cultured with AdMSCs at a 1:1 ratio. AdMSCs were used in passage 4. RPE-1 were also cultured alone for control. *The mixed cells (co-cultured cells)* were seeded at a density of 2 × 10^5^/cm^2^ per well in a plate, and cultured with an AdMSC medium supplemented with DMEM-LG with 10% FBS, 100 IU/ ml penicillin, 100 μg/ml streptomycin. The cells were incubated in a cell culture incubator containing 1% O2, for 48 hours at 37 ^o^C.

#### 2.4.2 Serum starvation model and in vitro co-culture system

RPE-1 with or without a *serum-free medium (serum starvation, SF)* were co-cultured with AdMSCs at a 1:1 ratio. The mixed cells were seeded at a density of 2 × 10^5^/cm^2^, and cultured with a serum-free medium supplemented with DMEM-LG with 100 IU/ ml penicillin and 100 μg/ml streptomycin. RPE-1 cells were also cultured alone. The cells were incubated at a cell culture incubator for 48 hours at 37 ^o^C.

### 2.5 Cell viability assay

CellTiter-Glo assay (CTG, Promega, G7570) was performed for cell viability. CTG reagent was mixed at a 1:10 ratio with supernatant from the treatment plate. The mix was incubated for 2 min at room temperature on the shaker and then allowed the plate to incubate at room temperature for 10 min to stabilize the luminescent signal, followed by luminescence measurement.

### 2.6 Determination of Reactive Oxygen Species (ROS)

RPE-1 with or without AdMSCs were seeded in 96 well plates and exposed to stress conditions for 48 hours. The cells were then incubated at 37 °C and 5% CO2 for 30 minutes with the 2,7-dichlorodihydrofluorescein diacetate (H_2_DCFH-DA) probe. At the end of the incubation, the medium was removed and replaced with the new medium. The fluorescence intensity was measured through a fluorescent Synergy-H1 Microplate Reader (Bio-Tek Instruments) with an excitation wavelength of 485 nm and an emission wavelength of 538 nm.

### 2.7 Microscopic Analysis

#### 2.7.1 Immunofluorescence (IF) Staining

Co-cultured cells and control groups were fixed with 4% PFA for 20 minutes at room temperature. Then, permeabilization of the cells was performed with 1% Triton X-100 (Sigma-Aldrich, T8787) in 1x DPBS for 7 minutes, and nonspecific antibody binding was blocked by incubating samples with a superblock solution (Scytek, AAA125) for 10 minutes. Then, cells were stained with primary antibodies and incubated for 90 minutes at 37 °C in a 1:50 dilution. To visualize the actin cytoskeleton, cells were incubated with monoclonal mouse anti-β-tubulin antibody (Abcam, ab6046), and negative control reactions were incubated with phosphate-buffered saline instead of the primary antibody. The secondary antibody, anti-mouse (1:100), was then used against the primary antibodies. Texas Red™-X Phalloidin (ThermoFisher, T7471) was used for binding F-actin of tunneling nanotubes and DAPI for nuclei.

#### 2.7.2 Quantification of TNTs

The numbers of TNTs in the IF-labelled populations were counted and expressed as the number of TNTs per 100 cells. Co-cultures of AdMSC and RPE-1 were incubated in stress. For counting of TNTs, AdMSC and RPE-1 were co-cultured for 48 hours in the presence of serum-free media. Afterward, cells were stained with anti-β-tubulin and Texas Red™-X Phalloidin and incubated for 90 minutes at 37 °C, followed by immunofluorescence imaging using the Leica TCSSP5 confocal microscope (TCS SP8 DLS, Leica) equipped with a 63x oil-immersion objective. 20 images containing at least 350 cells were obtained in each condition.

#### 2.7.3 Scanning electron microscopy (SEM) imaging

The media was discarded, and the cells were washed with DPBS. 2% PFA was prepared from 16% PFA solution (Paraformaldehyde 16% Solution, EM Grade, Electron Microscopy Sciences, Hatfield, PA) in nanopure water. The cells were fixed using a 2% PFA solution and let air dry for 2 days. SEM imaging was performed by ZEISS EVO LS15 Scanning electron microscope.

#### 2.7.4 Assessment of mitochondrial transfer

Briefly (Wittig, Wang et al. 2012), for live time-lapse imaging, co-cultured cells were monitored with a confocal microscope (TCS SP8 DLS, Leica) equipped with Differential Interference Contrast (DIC) components. Cells were imaged every 4 minutes at 37 °C and 5% CO2. Image analysis was performed using ImageJ/Fiji software (Version 2.0, PMID: 29187165). To stain mitochondria, cells were labeled with 1 mM JC-1 (Invitrogen) for 15 minutes at 37°C and visualized with a confocal microscope. JC-1 selectively accumulates within the mitochondrial matrix and forms red fluorescent J-aggregates (emission 590 nm) in the presence of negative transmembrane potential but exists as green monomers (emission 530 nm) under depolarised conditions. The green phase was recorded using an excitation wavelength of 485 nm and an emission filter of 540/550 nm. The red phase of JC-1 was recorded using an excitation wavelength of 535 nm and an emission filter of 610/675 nm.

### 2.8 Transmembrane (Transwell^®^) culture assay

Cell-cell contact (TNT Formation) is prevented by using two types of insert settings with a 0.4-μm pore polyester membrane (Transwell^®^, SARSTEDT) in a 24-well plate. AdMSCs were cultured on the upper surface of each insert membrane, and RPE-1 were seeded 24-well plate surface (Fig.5A), and vice versa (Fig5B). In the first set-up, AdMSCs/RPE-1 were allowed to co-culture for approximately 48 h, followed by removal of the insert and RPE-1 were analyzed for cell viability and ROS detection assays as described above.

### 2.9 Statistical analysis

Statistical analysis was performed using the Prism 8.02 (263) Software (GraphPad Software for Windows, San Diego, CA, USA), and results are reported as mean±S.D. For statistical analyses, the data were replicated in at least three experiments. Comparisons between more than two groups were analyzed by a one-way ANOVA test. Comparisons between the two groups were performed using Student’s t-test or two-way ANOVA test. Statistical significance was determined by P values less than 0.05.

## 3 Results

### 3.1 Cellular viability of RPE-1 cells in stress environments

Serum starvation (cultured in serum-free media) and oxidative stress (hypoxia) models induced TNTs. Cell survival was determined by CTG assay. The cell viability of RPE-1 seeded under stress without AdMSCs (RPE-1) was lower than that of the co-cultured (AdMSC/RPE-1) (Fig. 1A-B).

**Fig. 1.**
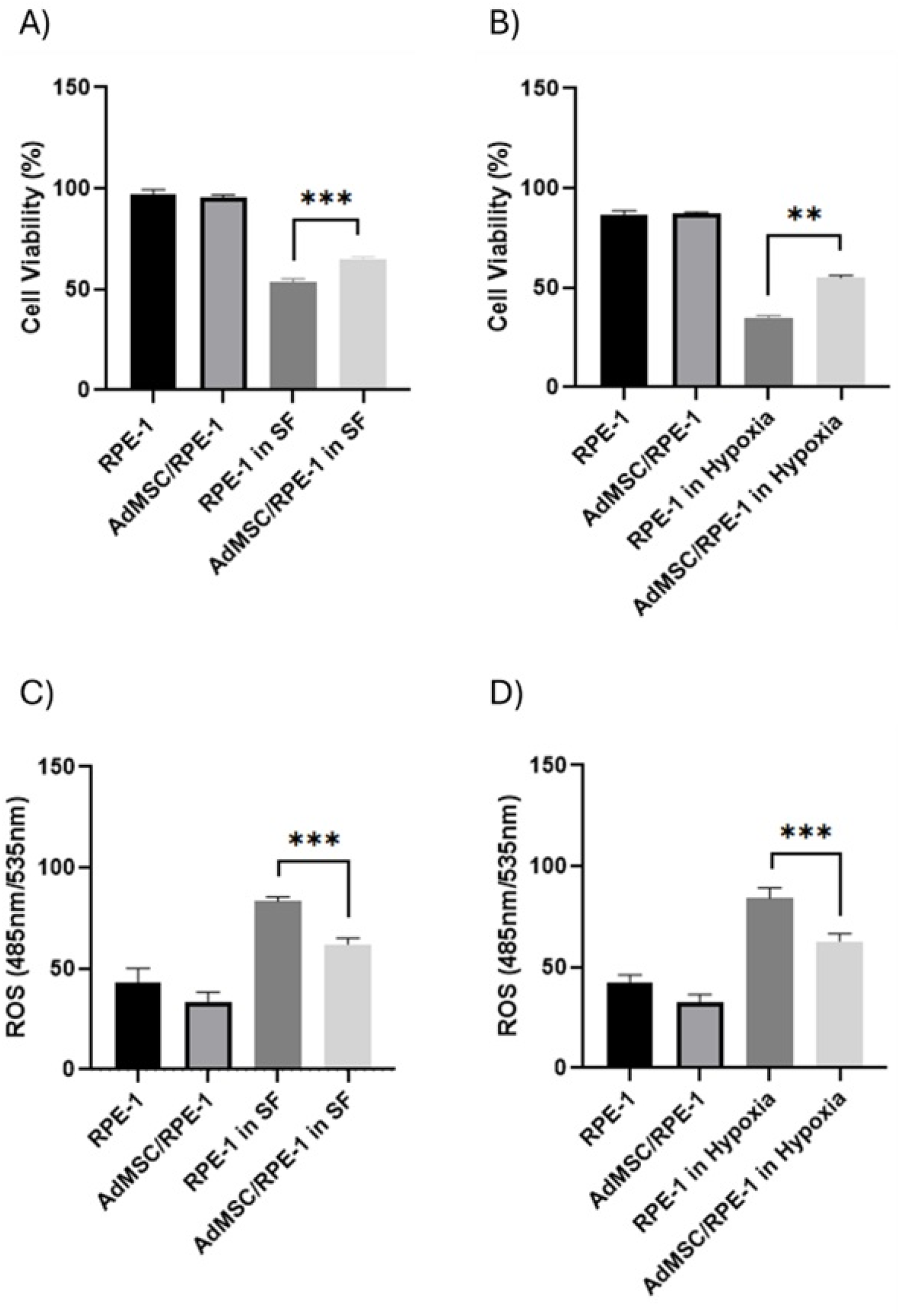
Cell viability and comparison of cells’ reactive oxygen species (ROS) in stress environments. **(A-B)** Cell viability assay was measured using CellTiter-Glo (CTG) at 48 hours. Cell viability significantly increased under stress conditions in co-cultured (AdMSC/RPE-1) compared to retina pigment epithelium (RPE)-1 seeded alone (RPE-1). **(C-D)** The change of ROS levels after 48 h of incubation with stress environments, where ROS levels significantly decreased under stress conditions in AdMSC/RPE-1 compared to RPE-1. SF: Serum Free (** P < 0.001 and *** P < 0.001).

### 3.2 Reactive oxygen species (ROS) levels in the co-culture group

Elevated intracellular ROS level causes macromolecule disruption, cell cycle arrest, and DNA damage, leading to the selective death of cells. 2,7-dichlorodihydrofluorescein diacetate (H_2_DCFH-DA) was employed to investigate the generation of ROS. In the presence of ROS, the H2DCF-DA molecule was converted to fluorescent dichlorofluorescein (DCF). The DCF fluorescence intensity was measured. Total ROS production in RPE-1 under stress conditions was significantly lower in the AdMSC/RPE-1 compared to RPE-1 (Fig. 1C-D).

### 3.3 Formation of TNTs between the AdMSC and RPE-1 cells

The immunofluorescence technique (IF) identified TNTs between the RPE-1 and AdMSCs. Co-cultured were found to be frequently interconnected by TNTs. Fluorescence microscopy revealed that TNTs, which are not anchored to the substratum, regularly connect cultured RPE-1 and AdMSCs in stress environments and contain F-actin but no microtubules (Fig. 2B-C). Scanning electron microscope (SEM) showed the ultrastructure of TNTs as straight connections between cells with a diameter ranging from 30 to 300 nm and a length of up to 120 mm (Supplemental Fig. 1).

**Fig. 2.**
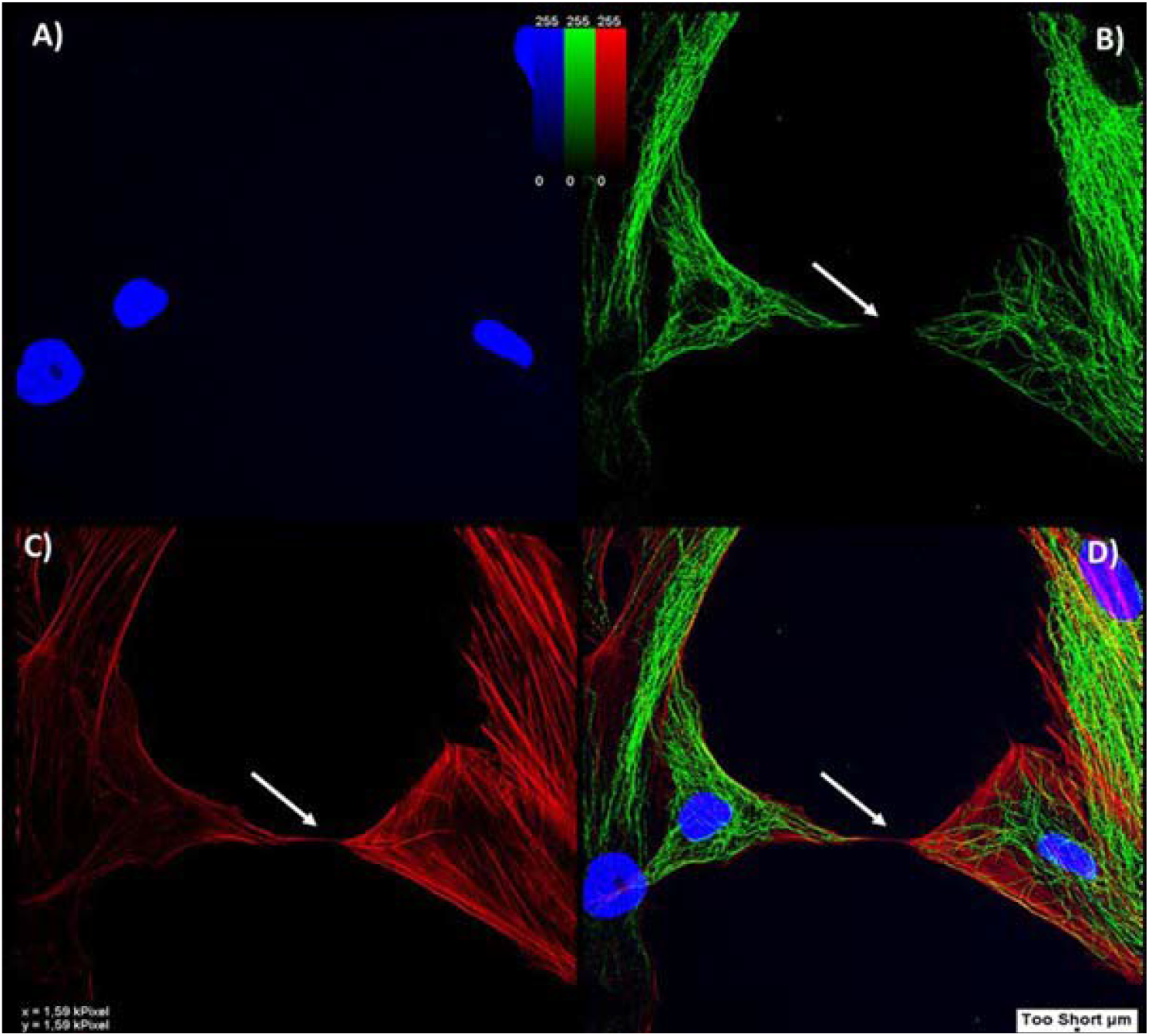
Tunneling Nanotubes (TNTs) form between the AdMSC and RPE-1. Immunofluorescence (IF) images of AdMSCs and RPE-1 stained with DAPI (blue) for nuclei **(A)**, anti-ß-tubulin (green) for microtubular structures **(B)**, and Texas Red™-X Phalloidin (red) for F-(actin) **(C). B-D:** IF images of two cells connected with TNT revealed the lack of ß-tubulin in the TNTs (arrows) and the presence of F-(actin) fibres within TNTs (arrows). **D:** Merged pictures with F- (actin) (red), ß-tubulin (green), and nucleus (blue).

### 3.4. Mitochondrial transfer from AdMSCs to RPE-1

The change in ROS levels in co-cultured cells suggests that the observed effects could be attributed to the transfer of mitochondria between the cells or the production of antioxidants by MSCs (Stavely and Nurgali 2020), for which we addressed TNTs formation between AdMSCs and RPE-1 in co-culture. By using the specific mitochondrial dye JC-1, we observed fluorescently-labelled-mitochondria inside TNTs of living cells and demonstrated that TNTs allowed mitochondria transfer from the AdMSCs to RPE-1 in serum starvation. (Fig. 3).

**Fig. 3.**
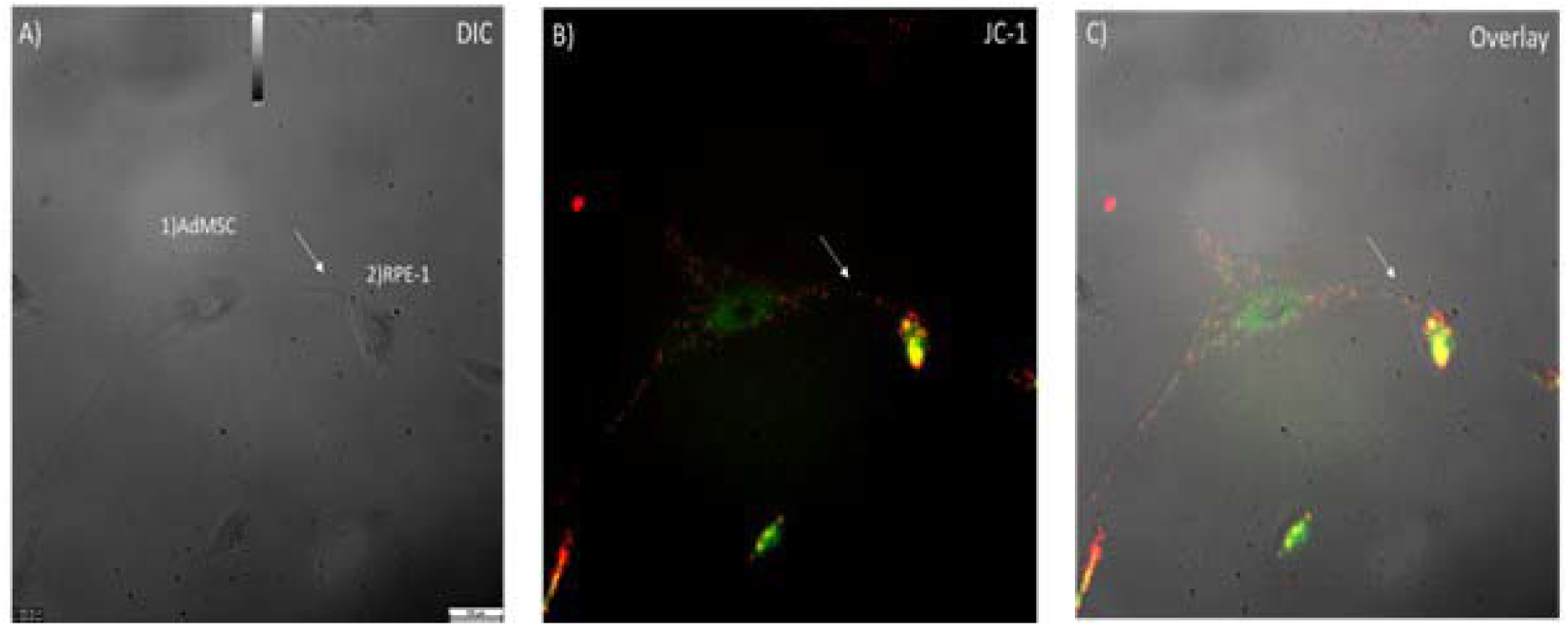
AdMSCs and RPE-1 are connected by tunneling nanotubes (TNT) containing mitochondria in serum starvation. **(A)** The bright field image shows two cells (1; AdMSC, 2; RPE-1) connected by a tunneling nanotube (arrow). The corresponding fluorescence image of **(A)** shows JC-1 labelled mitochondria of cells (arrow) **(B)**. The overlay of **(A)** and **(B)** shows the co-localization of nanotubes with fluorescent-labelled mitochondria **(C)**.

### 3.5 TNT formation increased in co-cultured cells compared to RPE-1, and it contributed to lower levels of ROS

To reveal the importance of TNT formation on the cells, we quantitatively analysed the number of TNTs between AdMSCs and RPE-1 for 48 hours. TNTs increased under serum starvation in both RPE-1 and co-cultured cells (AdMSC/RPE-1) (Fig. 4A). We have compared the fluctuation in ROS levels and the number of TNTs to interpret them in a time-dependent manner. When exposed to the stressful environment, cells in co-cultured were found more stress-resistant than the RPE-1. The stress response that started in the first 6 hours was reduced by 50% in the co-culture group at the end of 48 hours, while it was reduced by 45% in RPE-1 (Fig. 4B). Simultaneously, the TNT count had a more stable trend in the co-cultured. The initial number of co-cultured-TNTs was higher than the RPE-1 (Fig. 4C).

**Fig. 4.**
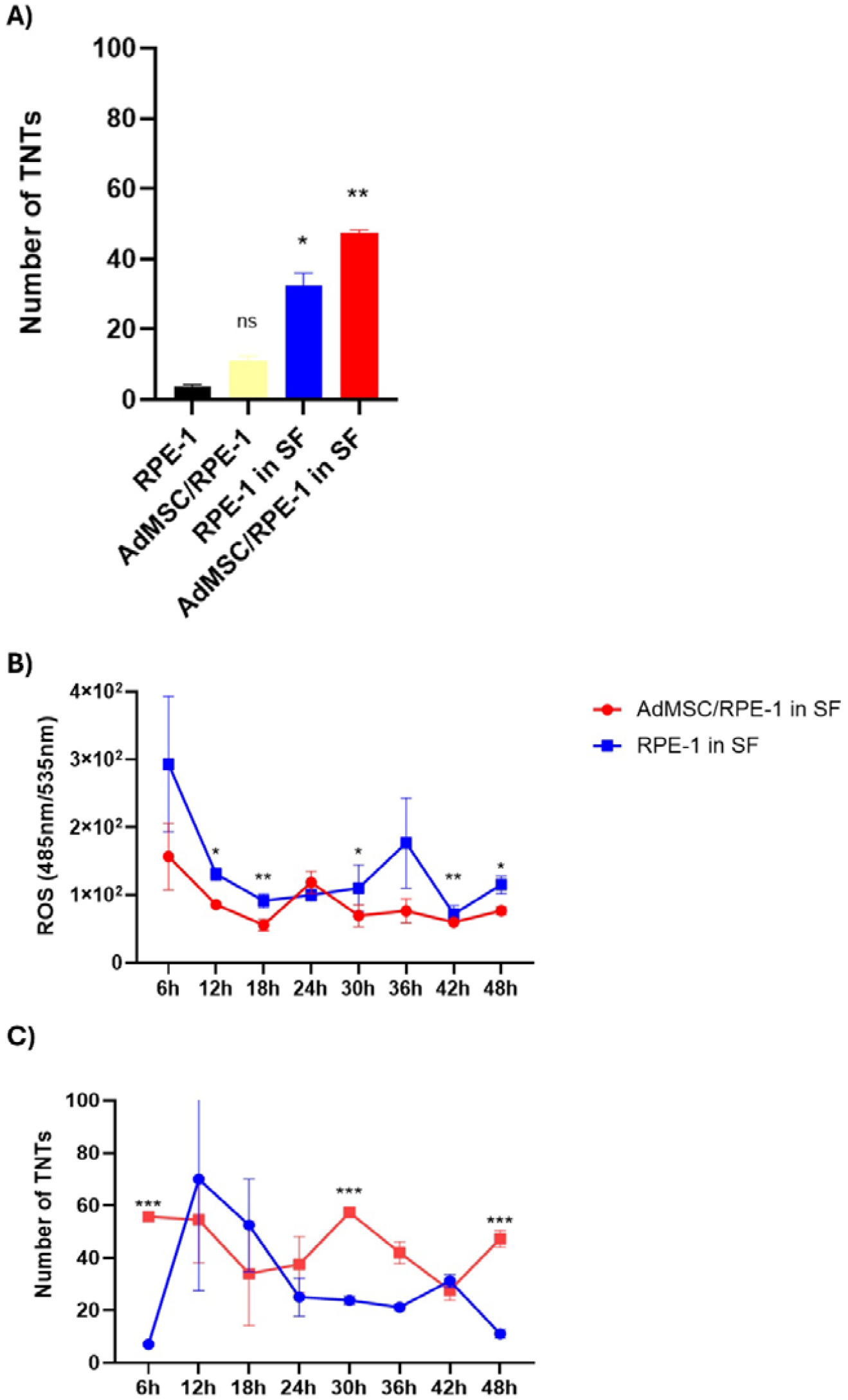
Effect of quantification of TNTs on cells in stress. **(A)** TNT structures were counted by immunofluorescence assay to show whether the survival of cells was related to TNTs. All groups were compared with RPE-1 in normal culture conditions. In the co-cultured, TNTs were more generated between RPE-1 and AdMSCs than RPE-1. **(B-C)** Time-dependent reactive oxygen species (ROS) levels **(B)** and the number of TNTs **(C)** were evaluated under serum starvation as a stress condition. Statistical significance was measured by comparing the groups. SF: Serum Free. (ns P>0.05,* P < 0.05, ** P < 0.01, ***P<0.001).

### 3.6 The Transwell^®^ culture system allowed TNT inhibition between the cells

Transwell^®^ culture systems (Fig. 5) managed to prevent TNT formation between AdMSCs and RPE-1. Flow cytometry detected the absence of mitochondrial transfer from JC-1 stained AdMSCs to RPE-1. The absence of transfer was interpreted as the absence of TNT formation (Fig. 5A-B). As no difference was observed between the two types of Transwell^®^ systems, further experiments were continued with the system shown in Fig. 7A to evaluate time-dependent cell viability and ROS changes in RPE-1. (Fig. 6B and 7A).

**Fig. 5.**
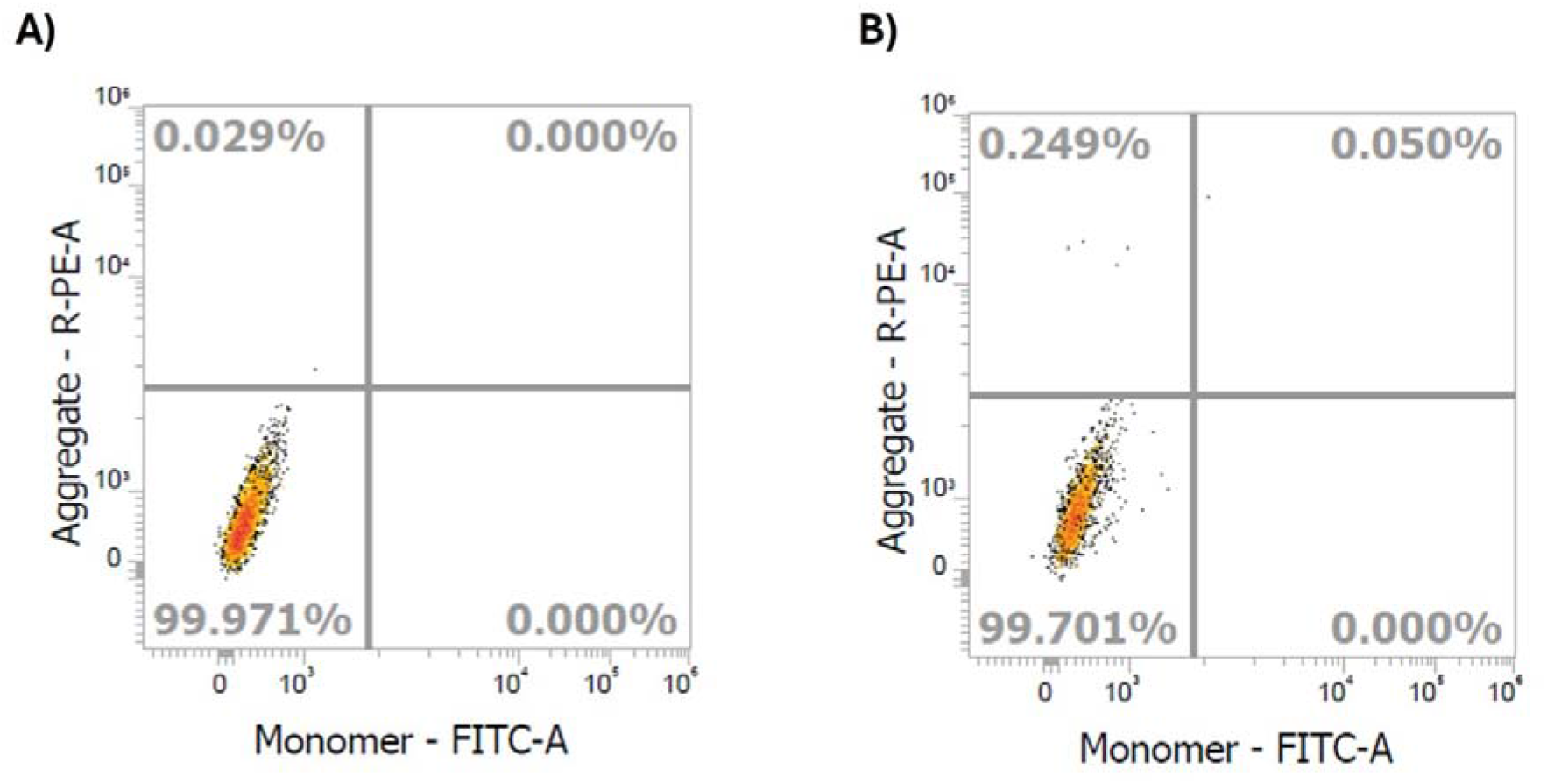
Schematic illustration of Transwell^®^ culture systems. Successful inhibition of intercellular TNTs under stress using an insert. Transwell^®^ membrane inserts (0.4 **μ**m pore size) were used to separate: A) the AdMSCs (insert) from RPE-1 (plate surface) and; B) the AdMSCs (plate surface) from RPE-1 cells (insert). AdMSCs were stained with JC-1 in both systems to analyze healthy mitochondria and displayed by flow cytometry. As no significant difference was observed between them, experiments were continued with the Transwell^®^ system depicted in **A** for better assessment of RPE-1 which are seeded on the plate.

**Fig. 6.**
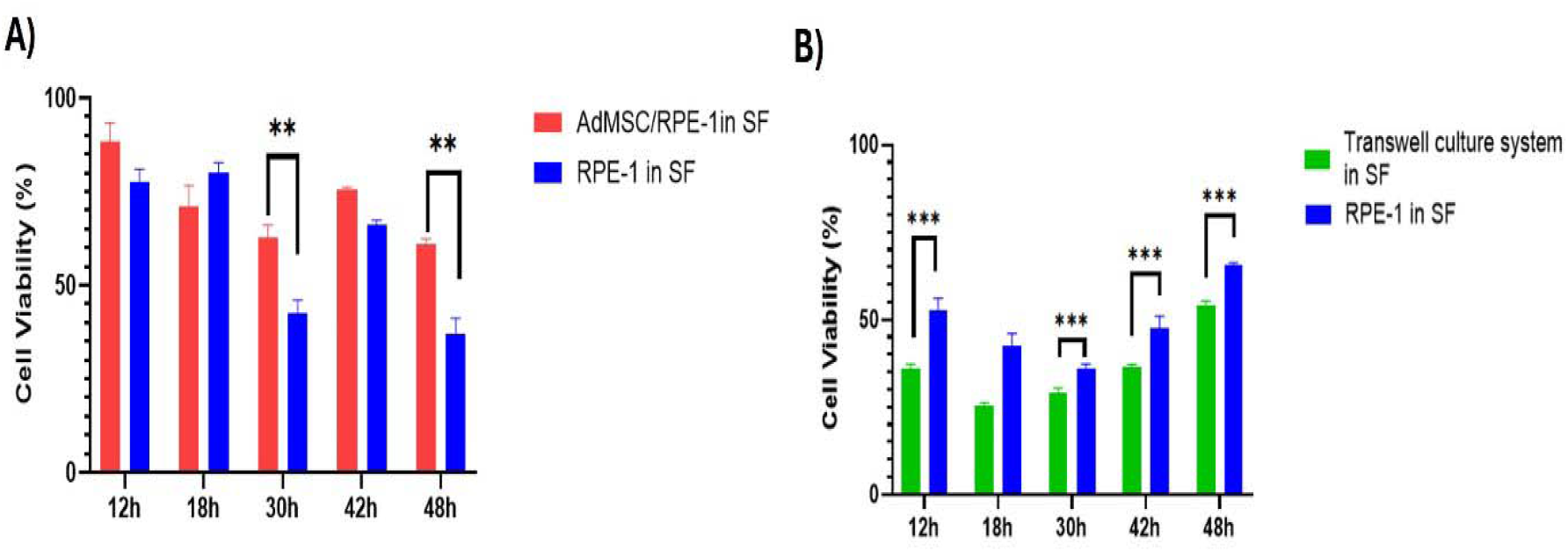
Comparison of the effects of co-cultured cells (A) and Transwell^®^ culture system (B) on RPE-1 viability. RPE-1 were seeded in a coculture or transwell system with AdMSCs. Cell viability assay was measured using CTG assay in a time-dependent manner. Statistical significance was measured by comparing the two groups. SF: Serum Free. (** P < 0.01 and *** P < 0.001).

**Fig. 7.**
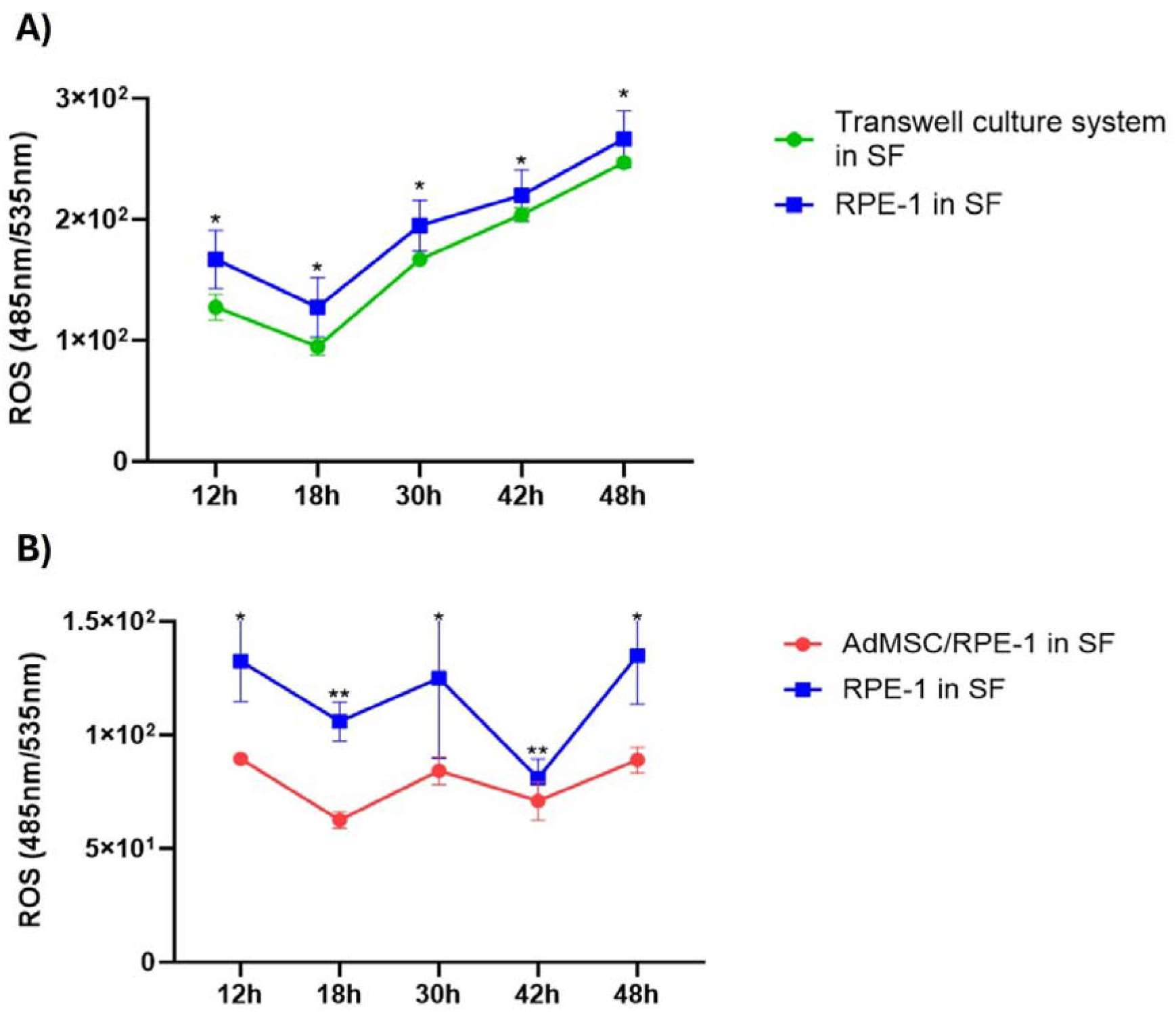
ROS changes of RPE-1 as time-dependently in the Transwell^®^ culture system and co-culture. Changes in ROS levels in RPE-1 in Transwell^®^ (A) and co-culture (B) were demonstrated in serum starvation, and ROS levels were not reduced when cells were separated by Transwell^®^ insert compared to co-culture. Statistical significance was measured by comparing between the two groups. SF: Serum Free. (* P < 0.05, ** P < 0.01)

### 3.7 Transwell^®^ culture system failed to improve cell viability and suppress ROS levels

Based on the results shown in Fig. 4B, cell viability and ROS levels of the RPE-1 of the Transwell^®^ culture system were analysed time-dependently by selecting the hours (12h, 18h, 30h, 42h, 48h) with statistically significant ROS levels. Under stress conditions, contrary to the co-culture system in which co-culture RPE-1 had higher viability scores than RPE-1, the RPE-1 cell viability was even found significantly lower than RPE-1 in Transwell^®^-separated cells (Fig. 6A-B).

To support the idea that AdMSCs need to contact RPE-1 in order to rescue retinal cells from stressful environments, we examined the time-dependent changes in ROS levels. Fig. 7A showed that the ability of retinal cells alone to cope with stress was still harder than those recruited in the Transwell^®^ culture system. Nevertheless, the Transwell^®^ culture system had a limited effect to reduce the stress of the cells for the first 18h, which remained insufficient afterwards. On the other hand, direct communication of cells in co-culture managed to keep ROS reduced when compared to baseline (Fig. 4B and 7B).

## Conclusions

Tunneling nanotubes (TNTs) are vital conduits for efficient cell-to-cell communication, facilitating the exchange of signals, chemicals, organelles, and pathogens among adjacent or distant cells. In the context of ocular tissues, including corneal, trabecular, and retinal cells, the presence of TNTs has been confirmed, underscoring their significance in ocular physiology and pathology. In the present study, we investigated the potential of adipose-derived mesenchymal stem cell (AdMSC) TNTs and AdMSC-TNTs-mediated mitochondrial transfer to determine how it helps in reducing reactive oxygen species (ROS) levels and preserving retinal pigment epithelium (RPE-1) viability under stress conditions. Our research includes intercellular mitochondrial transport from AdMSCs to RPE-1, revealing its importance, albeit not exclusively, in RPE-1 stress recovery via tunneling nanotubes (TNTs). Crucially, we demonstrated that direct interaction between AdMSCs and RPE-1 was essential for full stress recovery.

Mesenchymal cells (MSCs) have regenerative and immunomodulatory capabilities, which make them a promising treatment for a wide range of diseases, including autoimmune diseases, inflammatory airway disorders, liver diseases, muscle diseases, and neurodegenerative diseases (Fan, Zhang et al. 2020). Understanding the biology of MSCs and their role in treatment is critical to determine their potential for various therapeutic applications. Herein, we focussed on the relationship of AdMSC-therapies with TNTs. Our study determined the efficacy of co-culture conditions in enhancing viability and sustaining retinal pigment epithelial cell function after stress-induced damage (Fig. 1). The therapeutic potential of this co-culture technique in ocular tissue regeneration is evaluated by examining RPE cell survival. Studies on RGCs and CECs have shown that mitochondrial translocation via TNTs improves cell survival and tissue healing. Pro-inflammatory cytokines were found to be downregulated using quantitative cytokine array analysis, providing a foundation for investigating similar pathways in RPE protection (Jiang, Xiong et al. 2019). Afterward, we focused on examining the relationship between the quantity of TNTs and changes in ROS levels and cell viability. Increased nanotube numbers and reduced kidney tissue damage were observed in mouse kidney tissue under elevated oxidative stress, such as renal ischaemia or peritoneal dialysis (Ranzinger, Rustom et al. 2014). In a similar manner, the number of TNTs elevated in co-cultured cells after 48h serum starvation (Fig. 4A). MSCs can generate TNTs and deliver mitochondria and other components to target cells. This occurs under both normal and pathological situations, resulting in alterations in cellular energy metabolism and function (Luchetti, Carloni et al. 2022). Mitochondrial transfer successfully survived damaged cells afflicted with dysfunctional mitochondria by restoring their mitochondrial function, demonstrated through enhanced oxidative phosphorylation and increased ATP generation (Rustom 2016). Some studies showed that overexpression of calcium□sensitive adaptor proteins enhances the transfer of mitochondria from MSCs to neuron□like cells experiencing ischemic damage, improving recovery and cell proliferation (Ahmad, Mukherjee et al. 2014, Babenko, Silachev et al. 2018). Similarly, co-cultured mouse cardiomyocytes with human multipotent adipose-derived stem cells (hADSC) demonstrated that mitochondrial transfer from stem cells to cardiomyocytes plays a role in cardiomyocyte reprogramming (Acquistapace, Bru et al. 2011). By explaining intercellular mitochondrial trafficking from MSCs to corneal endothelial cells, photoreceptors, and retinal pigment cells, our study builds on earlier discoveries. According to recent studies, recipient cells that obtained mitochondria from MSCs demonstrate optimized respiratory capacities for the mitochondria and a greater expression of genes related to the structure and function of the mitochondria, suggesting the therapeutic potential of MSC-based mitochondrial transfer in the regeneration of ocular tissue (Jiang, Gao et al. 2016, Jiang, Chen et al. 2020). In particular, we discovered that AdMSCs may create TNTs that are directed toward target cells in order to transfer mitochondria, offering a direct channel for communication between cells and the donation of mitochondria (Fig. 3). Decreased total ROS production in RPE-1 under stress conditions was observed in co-culture (Fig. 4), supporting TNT-mediated mitochondrial transport. Since impaired mitochondria are a significant source of ROS due to disrupted oxidative phosphorylation, replacing them with functional mitochondria can effectively reduce ROS production. We confirmed that mitochondria passing from AdMSCs to unhealthy RPE-1 cells with increased TNT formation allowed ROS to decrease significantly and cell viability to increase (Fig. 4B-C); however, the unbalanced changes in ROS and TNT numbers demonstrated that mitochondria trafficking might not be the only factor. Cell survival of RPE-1 is also associated with other organelles such as ribosomes, Golgi vesicles, and macromolecules as nucleic acids and proteins (Sahu, Jena et al. 2018).

The ability of AdMSC-derived mitochondria to empower recipient RPE cells to cope with increased ROS levels is a central aspect of our study. By enhancing cellular antioxidant defence mechanisms and maintaining redox balance, we demonstrated the capacity of AdMSC-mediated mitochondrial transfer to mitigate oxidative stress-induced RPE damage (Fig. 4B, Supplemental Fig. 2). Similarly, studies on CECs have shown that mitochondrial transfer from AdMSCs reduces ROS levels and preserves mitochondrial function under oxidative stress, highlighting the therapeutic potential of this approach in protecting RPE against mitochondrial damage (Jiang, Gao et al. 2016). Therefore, indicating that intercellular mitochondrial transport as a vital mechanism for the regeneration of corneal epithelial cells and retinal ganglion cells, further supporting the role of mitochondrial transfer in ocular tissue repair. On the other hand, during 48 hours of serum deprivation, we observed the bidirectional mitochondrial exchange between MSCs and RPE-1 at various intervals (Supplemental Fig. 2). In vitro investigations revealed that mitochondrial transport can be bidirectional, which was previously demonstrated by the transfer of mitochondria between renal tubular cells and mesenchymal multipotent stromal cells (Plotnikov, Khryapenkova et al. 2010). Under normal culture conditions, bidirectional mitochondrial exchange was found between human vascular smooth muscle cells and BM-MSCs, which increased MSC proliferation (Dong, Rohlena et al. 2023). This exchange suggests a mutually beneficial effect on both cell populations in stress.

To show the effect of importance of TNTs formation, we forced a mechanical inhibition of TNTs via the transwell culture system (Fig. 5). Our findings support previous studies indicating that a transwell membrane filter can act as an efficient physical barrier to TNT formation (Thayanithy, O’Hare et al. 2017). After successfully inhibiting TNTs, we investigated the viability (Fig. 6) and ROS levels (Fig. 7) of RPE-1. When the cells were cultured together, they showed higher cell viability and a lower level of ROS than the transwell culture group. Moreover, RPE-1 viability decreased dramatically with the transwell culture system (Fig. 6). Indeed, several investigations have demonstrated the relevance of direct cell-cell contact in enabling metabolite transfer between cells, which can improve cell survival and function (Thayanithy, O’Hare et al. 2017). Therefore, we presume that a direct connection between the cells is essential. Furthermore, in co-culture systems or multicellular aggregates, physical contact between cells has been found to improve cell survival and functional outcomes as compared to cultures in which cells are spatially separated (Thayanithy, O’Hare et al. 2017). This shows that direct cell-to-cell interactions in stem cell treatment which allow TNT-mediated intercellular transfer could have advantages against indirect application of stem cell-based products such as exosomes (Harrell, Volarevic et al. 2022) for improving cellular responses to environmental stimuli and stresses in several tissue types as retinal pigment epithelium on the current applications. From the clinical perspective, it could be emphasized that the success of stem cell-based products on treatment of retinal diseases should also be dependent on the ability to form direct cell-to-cell connections between stem cells and target retinal cell groups according to different delivery methods including subretinal, intravitreal or suprachoroidal approaches (Battu, Ratra et al. 2022). Our findings highlight the importance of physical contacts between cells in facilitating intercellular communication and metabolic cooperation.

The main limitation of our study is the lack of comparing the effects of the secretome released specifically from adipose-derived stem cells with co-cultured and transwell assays, whose results could further elucidate the role of intercellular communication. In most of the studies, cell-free therapy seems to be favoured in regenerative medicine after the discovery that transplanted cells exert their therapeutic effects mainly through the secretion of paracrine factors (Foo, Looi et al. 2021). If the effects of the stem cell secretome are consistent with our findings, this shows that ‘the connection of cells’ is critical to the observed effects. This contrast can serve to support the concept that intercellular communication is both necessary and important in our experimental setups. By combining these various experimental methodologies, we can acquire a better knowledge of the mechanisms underpinning intercellular communication and its relevance in our study.

Our findings highlight the value of the direct connection of AdMSC and RPE-1 as a treatment method for shielding RPE against stress-induced damage. We also hope to emphasize the importance of the potential applications of AdMSC-mediated mitochondrial translocation via TNTs by exploring the mechanisms underpinning this method in retinal tissue repair and regeneration. Examples from RGC and CEC studies show that mitochondrial transfer via TNTs improves cell viability and promotes tissue regeneration through improved corneal wound healing after healthy MSC scaffold transplantation and transferred mitochondria detection in the corneal epithelium, supporting the hypothesis that similar mechanisms may contribute to RPE protection after retinal injury (Jiang, Gao et al. 2016).

## Supporting information

Supplemental Datas-updated

## Abbreviations

AdMSCs: Adipose-derived mesenchymal stem cells
MSCs: Mesenchymal stem cells
RPE-1: Retinal Pigment Epithelial Cell Line
TNTs: Tunneling nanotubes
ROS: Reactive oxygen species
DPBS: Dulbecco’s phosphate-buffered saline
DMEM-LG: Dulbecco’s Modified Eagle Medium - low glucose
P/S: penicillin/streptomycin
FBS: Fetal Bovine Serum
PBS: Phosphate-buffered saline
PFA: Paraformaldehyde
SF: Serum Free
H_2_DCFH-DA: 2,7-dichlorodihydrofluorescein diacetate
CTG: CellTiter-Glo
IF: Immunofluorescence
SEM: Scanning electron microscopy
DIC: Differential Interference
RGC: Retinal ganglion cells
CEC: Corneal epithelium cells

## Availability of data and materials

All data generated or analysed during this study are included in this article. The datasets used and/or analysed during the current study are available from the corresponding author on reasonable request.

## Author contributions

**Conceptualization;** Merve Gözel, **Data curation;** Merve Gözel, **Formal analysis;** Merve Gözel, **Funding acquisition;** Murat Hasanreisoğlu, **Investigation;** Merve Gözel, **Methodology;** Merve Gözel, **Project administration;** Merve Gözel, Murat Hasanreisoğlu, **Supervision;** Murat Hasanreisoğlu, Cem Kesim, **Validation;** Merve Gözel, **Visualization;** Merve Gözel **Roles/Writing - original draft;** Merve Gözel, Karya Senkoylu, Cem Kesim, Murat Hasanreisoğlu and **Writing - review & editing;** Merve Gözel, Karya Senkoylu, Cem Kesim, Murat Hasanreisoğlu

## Acknowledgments

The authors gratefully acknowledge the use of the services and facilities of the Koç University Research Center for Translational Medicine (KUTTAM) and Koç University Surface Science and Technology Center (KUYTAM), funded by the Presidency of Turkey, Head of Strategy and Budget. We sincerely thank Dr. Billur Sezgin for their invaluable assistance in providing the adipose tissue samples essential for this study. Her expertise was instrumental in the success of our research.

## References

Acquistapace, A., T. Bru, P. F. Lesault, F. Figeac, A. E. Coudert, O. le Coz, C. Christov, X. Baudin, F. Auber, R. Yiou, J. L. Dubois-Randé and A. M. Rodriguez (2011). “Human mesenchymal stem cells reprogram adult cardiomyocytes toward a progenitor-like state through partial cell fusion and mitochondria transfer.” Stem Cells 29(5): 812–824. DOI: 10.1002/stem.632

Ahmad, T., S. Mukherjee, B. Pattnaik, M. Kumar, S. Singh, M. Kumar, R. Rehman, B. K. Tiwari, K. A. Jha, A. P. Barhanpurkar, M. R. Wani, S. S. Roy, U. Mabalirajan, B. Ghosh and A. Agrawal (2014). “Miro1 regulates intercellular mitochondrial transport & enhances mesenchymal stem cell rescue efficacy.” Embo j 33(9): 994–1010. DOI: 10.1002/embj.201386030

Babenko, V. A., D. N. Silachev, V. A. Popkov, L. D. Zorova, I. B. Pevzner, E. Y. Plotnikov, G. T. Sukhikh and D. B. Zorov (2018). “Miro1 Enhances Mitochondria Transfer from Multipotent Mesenchymal Stem Cells (MMSC) to Neural Cells and Improves the Efficacy of Cell Recovery.” Molecules 23(3). DOI: 10.3390/molecules23030687

Battu, R., D. Ratra and L. Gopal (2022). “Newer therapeutic options for inherited retinal diseases: Gene and cell replacement therapy.” Indian J Ophthalmol 70(7): 2316–2325. DOI: 10.4103/ijo.IJO_82_22

Chinnery, H. R. and K. E. Keller (2020). “Tunneling Nanotubes and the Eye: Intercellular Communication and Implications for Ocular Health and Disease.” Biomed Res Int 2020: 7246785. DOI: 10.1155/2020/7246785

Dong, L. F., J. Rohlena, R. Zobalova, Z. Nahacka, A. M. Rodriguez, M. V. Berridge and J. Neuzil (2023). “Mitochondria on the move: Horizontal mitochondrial transfer in disease and health.” J Cell Biol 222(3). DOI: 10.1083/jcb.202211044

Dupont, M., S. Souriant, G. Lugo-Villarino, I. Maridonneau-Parini and C. Vérollet (2018). “Tunneling Nanotubes: Intimate Communication between Myeloid Cells.” Front Immunol 9: 43. 10.3389/fimmu.2018.00043

Eiro, N., J. Sendon-Lago, S. Cid, J. Saa, N. de Pablo, B. Vega, M. A. Bermudez, R. Perez-Fernandez and F. J. Vizoso (2022). “Conditioned Medium from Human Uterine Cervical Stem Cells Regulates Oxidative Stress and Angiogenesis of Retinal Pigment Epithelial Cells.” Ophthalmic Res 65(5): 556–565. DOI:10.1159/000524484

Fan, X. L., Y. Zhang, X. Li and Q. L. Fu (2020). “Mechanisms underlying the protective effects of mesenchymal stem cell-based therapy.” Cell Mol Life Sci 77(14): 2771–2794. DOI: 10.1007/s00018-020-03454-6

Foo, J. B., Q. H. Looi, P. P. Chong, N. H. Hassan, G. E. C. Yeo, C. Y. Ng, B. Koh, C. W. How, S. H. Lee and J. X. Law (2021). “Comparing the Therapeutic Potential of Stem Cells and their Secretory Products in Regenerative Medicine.” Stem Cells Int 2021: 2616807. DOI: 10.1155/2021/2616807

Gozel, M., et al., Effects of Primed Adipose Mesenchymal Stem Cell-Derived Exosomes on Immunomodulation in Behcet Uveitis. bioRxiv, 2024: p. 2024.11.24.625042.

Harrell, C. R., V. Volarevic, V. Djonov and A. Volarevic (2022). “Therapeutic Potential of Exosomes Derived from Adipose Tissue-Sourced Mesenchymal Stem Cells in the Treatment of Neural and Retinal Diseases.” Int J Mol Sci 23(9). DOI: 10.3390/ijms23094487

Jiang, D., F. X. Chen, H. Zhou, Y. Y. Lu, H. Tan, S. J. Yu, J. Yuan, H. Liu, W. Meng and Z. B. Jin (2020). “Bioenergetic Crosstalk between Mesenchymal Stem Cells and various Ocular Cells through the intercellular trafficking of Mitochondria.” Theranostics 10(16): 7260–7272. DOI:10.7150/thno.46332

Jiang, D., F. Gao, Y. Zhang, D. S. Wong, Q. Li, H. F. Tse, G. Xu, Z. Yu and Q. Lian (2016). “Mitochondrial transfer of mesenchymal stem cells effectively protects corneal epithelial cells from mitochondrial damage.” Cell Death Dis 7(11): e2467. DOI: 10.1038/cddis.2016.358

Jiang, D., G. Xiong, H. Feng, Z. Zhang, P. Chen, B. Yan, L. Chen, K. Gandhervin, C. Ma, C. Li, S. Han, Y. Zhang, C. Liao, T. L. Lee, H. F. Tse, Q. L. Fu, K. Chiu and Q. Lian (2019). “Donation of mitochondria by iPSC-derived mesenchymal stem cells protects retinal ganglion cells against mitochondrial complex I defect-induced degeneration.” Theranostics 9(8): 2395–2410. DOI:10.7150/thno.29422

Luchetti, F., S. Carloni, M. G. Nasoni, R. J. Reiter and W. Balduini (2022). “Tunneling nanotubes and mesenchymal stem cells: New insights into the role of melatonin in neuronal recovery.” J Pineal Res 73(1): e12800. DOI: 10.1111/jpi.12800

Pittenger, M. F., A. M. Mackay, S. C. Beck, R. K. Jaiswal, R. Douglas, J. D. Mosca, M. A. Moorman, D. W. Simonetti, S. Craig and D. R. Marshak (1999). “Multilineage potential of adult human mesenchymal stem cells.” Science 284(5411): 143–147. DOI: 10.1126/science.284.5411.143

Shi, Y., et al., Advancements in culture technology of adipose-derived stromal/stem cells: implications for diabetes and its complications. Front Endocrinol (Lausanne), 2024. 15: p.1343255.

Stavely, R. and K. Nurgali, The emerging antioxidant paradigm of mesenchymal stem cell therapy. Stem Cells Transl Med, 2020. 9(9): p. 985–1006.

Plotnikov, E. Y., T. G. Khryapenkova, S. I. Galkina, G. T. Sukhikh and D. B. Zorov (2010). “Cytoplasm and organelle transfer between mesenchymal multipotent stromal cells and renal tubular cells in co-culture.” Exp Cell Res 316(15): 2447–2455. DOI: 10.1016/j.yexcr.2010.06.009

Quigley, H. A. (2011). “Glaucoma.” Lancet 377(9774): 1367–1377. DOI: 10.1016/S0140-6736(10)61423-7

Ranzinger, J., A. Rustom, D. Heide, C. Morath, P. Schemmer, P. P. Nawroth, M. Zeier and V. Schwenger (2014). “The receptor for advanced glycation end-products (RAGE) plays a key role in the formation of nanotubes (NTs) between peritoneal mesothelial cells and in murine kidneys.” Cell Tissue Res 357(3): 667–679. DOI: 10.1007/s00441-014-1904-y

Rustom, A. (2016). “The missing link: does tunnelling nanotube-based supercellularity provide a new understanding of chronic and lifestyle diseases?” Open Biol 6(6). 10.1098/rsob.160057

Sahu, P., S. R. Jena and L. Samanta (2018). “Tunneling Nanotubes: A Versatile Target for Cancer Therapy.” Curr Cancer Drug Targets 18(6): 514–521. DOI: 10.2174/1568009618666171129222637

Thayanithy, V., P. O’Hare, P. Wong, X. Zhao, C. J. Steer, S. Subramanian and E. Lou (2017). “A transwell assay that excludes exosomes for assessment of tunneling nanotube-mediated intercellular communication.” Cell Commun Signal 15(1): 46. DOI: 10.1186/s12964-017-0201-2

Tiwari, V., R. Koganti, G. Russell, A. Sharma and D. Shukla (2021). “Role of Tunneling Nanotubes in Viral Infection, Neurodegenerative Disease, and Cancer.” Front Immunol 12: 680891. 10.3389/fimmu.2021.680891

Wittig, D., X. Wang, C. Walter, H. H. Gerdes, R. H. Funk and C. Roehlecke (2012). “Multi-level communication of human retinal pigment epithelial cells via tunneling nanotubes.” PLoS One 7(3): e33195. DOI: 10.1371/journal.pone.0033195

Yang, S., J. Zhou and D. Li (2021). “Functions and Diseases of the Retinal Pigment Epithelium.” Front Pharmacol 12: 727870. DOI: 10.3389/fphar.2021.727870

Zhu, X., Z. Chen, L. Wang, Q. Ou, Z. Feng, H. Xiao, Q. Shen, Y. Li, C. Jin, J. Y. Xu, F. Gao, J. Wang, J. Zhang, J. Zhang, Z. Xu, G. T. Xu, L. Lu and H. Tian (2022). “Direct conversion of human umbilical cord mesenchymal stem cells into retinal pigment epithelial cells for treatment of retinal degeneration.” Cell Death Dis 13(9): 785. 10.1038/s41419-022-05199-5

