## Supplemental Datas-updated for "Adipose-derived Mesenchymal Stem Cells and Retinal Pigment Epithelial Cells Interactions in Stress Environment via Tunneling Nanotubes"

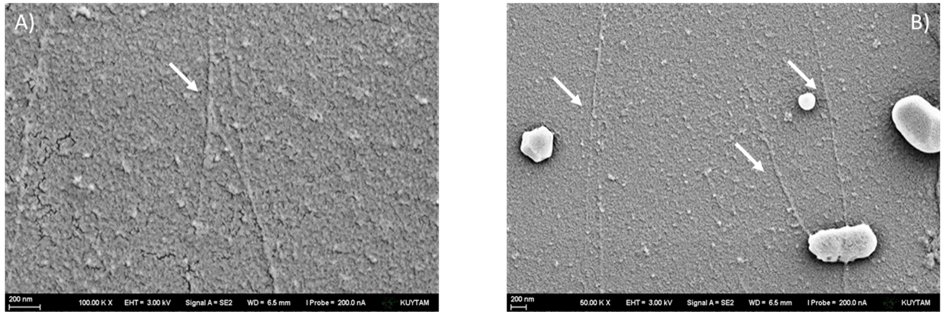

**Supplementary Fig. 1.** The presence of tunneling nanotubes (TNTs) in AdMSC and RPE-1 cells by scanning electron microscopy (SEM). TNTs form in (A) serum starvation and (B) a hypoxic environment. (A, B) SEM imaging shows the ultrastructure of TNTs between the AdMSCs and RPE-1 cells. TNTs are straight connections between two cells and have typical diameters of 30-300 nm. Arrows mark two focal thickenings of 1-2 mm, indicating a possible transport of organelles through the tunneling nanotubes. The picture was taken with 3.00 kV. Scale bar, 200 nm. AdMSC: Adipose derived mesenchymal stem cell. RPE: retina pigment epithelium.

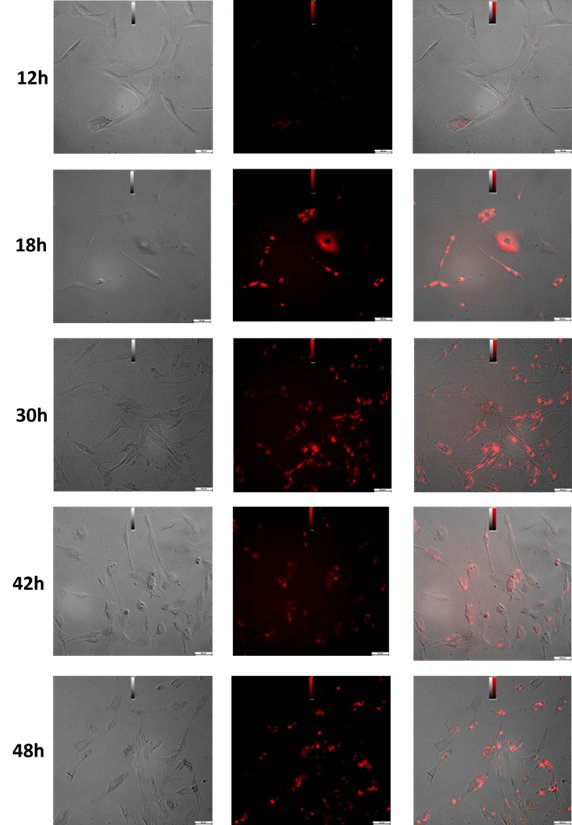

**Supplementary Fig. 2.** **Time-dependent changes in the number of TNTs and ROS levels and evaluation of significant points by mitochondrial transitions.** The bright field image shows two cells connected by a tunneling nanotube**.** The corresponding fluorescence image shows JC-1 labeled mitochondria of cells**.** The overlay shows the co-localization of nanotubes with fluorescent-labeled mitochondria in time-dependently**.**
